# FLP-15 functions through the GPCR NPR-3 to regulate local and global search behaviours in *Caenorhabditis elegans*

**DOI:** 10.1101/2025.05.02.651881

**Authors:** Umer Saleem Bhat, Siju Surendran, Sharanya H, Jun Liu, Yun Xu, Namra Tasnim, Ashwani Bhardwaj, Monika Scholz, Kavita Babu

**Affiliations:** Centre for Neuroscience, Indian Institute of Science (IISc), Bangalore, Karnataka, India; Department of Biological Sciences, Indian Institute of Science Education and Research (IISER) Mohali, Sector 81, SAS Nagar, Manauli, Punjab, India; Max Planck Research Group Neural Information Flow, Max Planck Institute for Neurobiology of Behavior – Caesar, Bonn, Germany

**Author notes:** Université de Fribourg, Switzerland (Siju Surendran). BRIC-National Institute of Immunology, New Delhi, India.

**Keywords:** FLP-15, NPR-3, *C. elegans*, local search, global search

## Abstract

Foraging is essential for sustenance and well-being of all organisms. The transition from well-fed to food-deprived conditions in *C. elegans* triggers a localized exploration of the environment characterized by frequent reorientations. However, over time the cumulative frequency of these reorientations decreases, facilitating the transition to global search behaviour. To investigate the genetic regulation of foraging in *C. elegans*, we conducted a screen of neuropeptide mutants and identified several candidates involved in modulating this behaviour. Among these, neuropeptide FLP-15 emerged as a key regulator of both local and global search behaviours. Our observations revealed that FLP-15 regulates the frequency and duration of reversals during foraging. Further investigation indicated that FLP-15 is expressed in and functions through the I2 pharyngeal neuron via the G-protein coupled receptor NPR-3. Mutants lacking either *flp-15* or *npr-3* displayed a significant decrease in reversal frequency during local search behaviours. Interestingly, unlike wild-type animals, the reversal frequency in *flp-15* and *npr-3* mutants did not decrease over time. This study also describes the expression pattern of NPR-3, in a subset of head neurons, predominantly comprising of dopaminergic neurons. This expression pattern highlights a potential link between neuropeptide signalling and dopaminergic modulation of behaviour. Finally, exogenous dopamine supplementation assays revealed that FLP-15 may regulate foraging by modulating dopamine transmission, highlighting a novel neuropeptide-dopamine interaction involved in the control of foraging behaviours.

## Introduction

Neurons communicate with each other through small neurotransmitters in chemical synapses and through gap junctions in electrical synapses (reviewed in (Borroto-Escuela *et al*, 2024; Sheffler *et al*, 2025)). However, extrasynaptic communication, extending beyond synapses, is facilitated by another class of molecules known as neuropeptides (Li *et al*, 2020; Xiong *et al*, 2023). A notable distinction in the action of neuropeptides compared to other forms of neuronal communication is the slow and sustained nature of neuropeptide signalling (reviewed in (van den Pol, 2012)). This temporal profile arises from the diffusion of neuropeptides, subsequent to their release, through volumetric transmission, and also due to the absence of neuropeptide reuptake machinery in neurons (reviewed in (van den Pol, 2012)). Furthermore, neuropeptides signal through G-protein coupled receptors (GPCR) at time-scales slower than neurotransmitters functioning through ionic receptors, hence enabling prolonged effects (reviewed in (Guillaumin & Burdakov, 2021; van den Pol, 2012)).

Neuropeptides have been reported to play critical roles in modulating a wide range of physiological processes across phyla including locomotion, appetite, mood, pain perception, reproduction, sleep, and learning and memory (reviewed in (Bhat *et al*, 2021; Carniglia *et al*, 2017)). The stereotypic locomotion of *C. elegans* is characterized by distinct patterns, encompassing forward movement interspersed with reversals and omega turns (Gray *et al*, 2005). Despite numerous studies aimed at elucidating the intricate neuronal circuitry that controls each aspect of *C. elegans* locomotion, the underlying mechanism and molecular players involved are still largely unknown (Gray *et al*., 2005; Piggott *et al*, 2011). Recent work has parsed out the neuropeptidergic connectome in *C. elegans* with neuropeptides emerging as important signalling molecules that play a crucial regulatory role in modulating multiple locomotory characteristics ((Ripoll-Sanchez *et al*, 2023) and reviewed in (Venkatesh *et al*, 2025; Watteyne *et al*, 2024)).

Neuropeptides have been implicated in modulating sustained responses, across phyla. Starvation and satiety are some of the most widely studied sustained responses as they are vital aspects of nutrition in all organisms. In most vertebrates, an orexigenic neuropeptide such as Ghrelin is secreted from the stomach during hunger and starvation whereas anorexigenic neuropeptides such as cholecystokinin, glucagon like peptide (GLP-1), insulin, and leptin, pancreatic polypeptide (PP), and PYY are secreted in response to satiety (reviewed in (Alhabeeb *et al*, 2021; Wisser *et al*, 2010)). These neuropeptides serve as indicators of the internal metabolic state of the organism and further modulate the locomotor circuit to execute foraging strategies like dwelling, food seeking and roaming. Although defects in these peptidergic pathways could predispose an organism to metabolic syndromes, the literature in this area remains scant.

In this study we have used food-related behavioural states to delve into the mechanism underlying neuromodulation of sustained behaviours by neuropeptides, specifically, foraging in *C. elegans*. When *C. elegans* are transferred from well-fed conditions to a nutrient deprived state, they initially explore locally with a combination of reorientations (reversals and omega turns) executed frequently. This search behaviour is followed by a decrease in the cumulative frequency of reorientations to give rise to a global search of the arena (Lopez-Cruz *et al*, 2019). Defects in reorientations and/or the body wave parameters like wavelength and amplitude result in inefficient exploration of the environment (Oranth *et al*, 2018; Ramachandran *et al*, 2021).

We hypothesize that neuropeptides play a critical role in integrating metabolic and sensory cues with behavioural responses, allowing organisms to adjust their foraging behaviour. Therefore, in this study, our primary objective was to elucidate the role of neuropeptides as molecular indicators of the metabolic status of the organism and examine how these signalling molecules influence the animal’s exploratory strategy during foraging.

## Materials and Methods

### Strains maintenance

All strains were cultured on nematode growth medium (NGM) plates, seeded with OP50 *E. coli* bacteria, and maintained under standard conditions at 22°C for most experiments and 20°C for the pharyngeal pumping and speed analyses. The strains were maintained following established protocols outlined in previous literature (Brenner, 1974). The N2 Bristol strain served as the standard wild-type (WT) control for comparative analyses across all experimental procedures. A synchronous population of young adult *C. elegans* was obtained using previously established protocol (Porta-de-la-Riva *et al*, 2012). The list of strains used in this study is tabulated in Supplementary Table 1 (S1).

### Molecular biology and transgenes

Transcriptional reporters were generated through molecular cloning of the promoter regions, positioning them upstream of fluorescent proteins (GFP) in the pPD95.75 vector or mCherry in the pPD49.26 vector. The length of all promoter sequences was maintained at 2 kb, spanning the genomic region upstream of the start codon for each respective gene and the sequences were extracted from WormBase. Promoters of genes *flp-15, gur-3, npr-3, dat-1 and nlp-12* have been used in this study. The rescue constructs were generated by cloning genomic DNA or cDNA downstream of the respective promoter. All the constructs were microinjected in *C. elegans* to prepare transgenic lines as previously described (Mello & Fire, 1995; Mello *et al*, 1991). The list of primers and plasmids used in this study is tabulated in Supplementary Tables 2 and 3 (S2 and S3) respectively.

### CRISPR Cas9 genome editing

The *npr-3* null mutant was generated using the CRISPR Cas9 genome editing system. Two single-guide RNAs (sgRNAs) were designed using CRISPOR (Concordet & Haeussler, 2018), targeting binding sites in exon 1 and exon 6. The specific sequences for the sgRNAs, obtained from Integrated DNA Technologies (IDT), were TAGCTCCAACCCAAACAAGA for exon 1 and ATCAGTACCCTTGAACAAAG for exon 6. A reaction mixture composed of 0.25 µg/µL Cas9 enzyme (IDT), 2.5 µM of each sgRNA, and TE buffer (pH 7.5) to achieve a total volume of 5 µL, was incubated at 37°C for 15 minutes. Subsequently, 5 ng/µL of pCFJ90 plasmid (injection marker) was added to the reaction, and nuclease-free water was used to adjust the volume to 20 µL. This final mixture was used for microinjecting *C. elegans*, to generate the *npr-3* knockout. Following microinjection, 20-30 F1 *C. elegans* expressing the pharyngeal marker (pCFJ90) were singled out and allowed to produce progeny. The parent animals were then subjected to genotyping (Primers listed in Table S2) for the *npr-3* mutation. Heterozygous *C. elegans* with the 2799 bp deletion were homozygosed. The *npr-3* mutants then underwent three rounds of outcrossing with wild-type *C. elegans* to eliminate any non-specific background mutations.

### Single animal tracking during local and global search

All the single animal tracking assays were performed using single young adult *C. elegans* recorded one at a time across several days to ensure consistency and account for variability. Briefly, a well-fed young adult *C. elegans* was picked with an eyelash pick by using a small drop of halocarbon oil, ensuring a gentle transfer. The *C. elegans* was then placed in an off food NGM plate and allowed to crawl freely for two minutes to get rid of any residual food. The animal was then gently picked again and placed on the freshly prepared 90 mm NGM (without cholesterol) plate calibrated at room temperature and allowed to move freely for one minute before starting the tracking. The recording of *C. elegans* locomotion for screen purpose was done using 1.6 MP CMOS camera by Thor lab mounted on a Zeiss Stemi 2000-C stereoscope. Subsequent recordings were done using the MBF Bioscience WormLab imaging system at 8 fps. The C. elegans were recorded initially for a period of 5 minutes in off food condition (local search). Following the initial recording, the *C. elegans* was then allowed to explore the same plate for 25 minutes and then recorded again for 5 minutes (global search) under similar conditions (Illustrated in Figure S2A). For on food assay, a single well-fed *C. elegans* was transferred to a freshly and uniformly seeded OP50 plate and allowed to move freely for 2 minutes. The locomotion was then recorded for 5 minutes for further analysis. Frequency of reversals was determined by the total number of spontaneous reversals executed in five minutes. Reversals longer than two head swings were considered for the quantification of frequency. Reversal length was calculated based on the number of head swings performed during each spontaneous reversal, as previously described (Bhardwaj *et al*, 2018). The reversal length was calculated for each reversal and averaged over the total number of reversals made over five minutes which was then plotted as single data point. The number of *C. elegans* recorded for each genotype was ≥ 20.

### Multiple animal tracking for analysing pumping rate of *C. elegans*

Assays were performed on 15 cm imaging NGM plates (NGM without cholesterol and agarose replacing agar). Animals at the L4 stage were selected 12 hours before experiments for age synchronization. Young adults animals were transferred to the pre-prepared NGM plates using an eyelash pick and halocarbon oil. Following a 1 minute acclimation and starvation period under blue light, animals were recorded for 1 minute using a custom epi-fluorescence tracking microscope with a 15 ms exposure and a sampling rate of 30 frames/s. The microscope was configured with a white-light LED, and a GFP filter set. Global search assays were performed 25 minutes after the corresponding local search assay using the same plate with the same settings. Data was analysed with Code adapted from PharaGlow (Bonnard *et al*, 2022) and the mean pumping rate was extracted per animal.

### Exogenous dopamine supplementation assay

The exogenous dopamine supplementation assay was performed using 90 mm NGM plates supplemented with dopamine hydrochloride solution. To prepare the assay plates, 80 µL of freshly prepared 1 M dopamine hydrochloride solution (Sigma) was evenly spread onto the surface of each plate to achieve a final concentration of 40 mM, as previously described (Baidya *et al*, 2014; Pandey *et al*, 2021). The plates were allowed to dry in the dark for at least 10 minutes and were used immediately after. Behavioural recordings were carried out as described in the single *C. elegans* tracking protocol during the local search behaviour.

### Imaging experiments

All the gene expression studies were done using transgenic reporter lines carrying either GFP or mCherry as indicated in the Table S1. Young adult *C. elegans* were transferred to 2 μl of 2,3-butanedione monoxime (30 mg/ml) placed on a 2% agar pad and imaged after undergoing paralysis and being covered with a glass cover slip as previously described (Tikiyani *et al*, 2018). Images were acquired using an Andor Benchtop Confocal Microscope - BC43.

### Statistical analyses

The data was plotted on GraphPad Prism 10 and represented as violin or box and whisker plots to represent distribution of data. Error bars represent SEM. One-way ANOVA with multiple comparisons and post-hoc Tukey tests was performed to compare the results in most of the experiments. For calculating the pharyngeal pumps, significance was assessed using a Mann-Whitney U-test with a Holm-Sidak correction for multiple testing. The level of significance was set as p≤ 0.05.

## Results

### Neuropeptides regulate the reversal frequency during foraging

We aimed to dissect the neuropeptidergic modulation of locomotion during foraging in *C. elegans*. Therefore, we conducted a comprehensive screen of mutants in 22 neuropeptide candidate genes, each of which was expressed in neurons previously implicated in feeding or foraging circuits based on existing literature and databases (Gray *et al*., 2005; Piggott *et al*., 2011). During screening, we recorded the movement of each mutant strain for 5 minutes during local search (0-5 minutes) and another 5 minutes during the global search (25-30 minutes) as illustrated (Figure S1A). We specifically focused on analysing the frequency and length of reversals that are key aspects of locomotory behaviour associated with foraging and compared these parameters with those of the wild-type (WT) control strain (Figures 1A, B and S1B).

**Figure 1:**
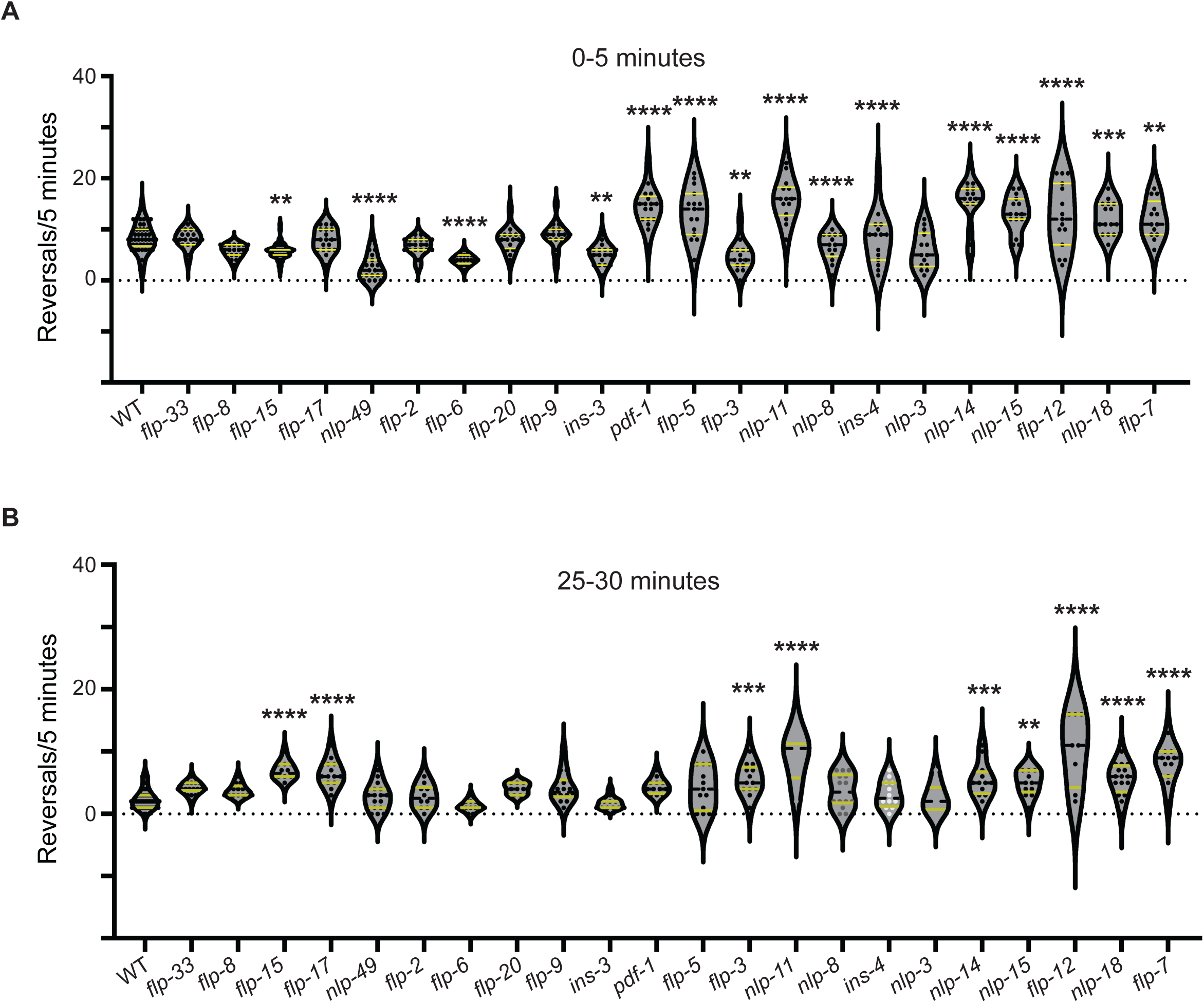
Neuropeptides regulate foraging by regulating reversal frequency during local and global search. (A-B) Violin plots showing quantification of reversals during local and global search respectively in neuropeptide mutants. Many of the neuropeptides screened showed a significant change in the frequency of reversals during local search in comparison to the WT. In this plot, each dot corresponds to the number of spontaneous reversals observed per 5 minutes from a single *C. elegans*. Statistical significance was determined using One-way ANOVA with Tukey test for multiple comparisons, “**”, “***’’ or “****” indicate p-values less than 0.01, 0.001 or 0.0001, respectively. The absence of “*” indicates not significantly different from WT control animals.

Our results revealed that many neuropeptide mutants exhibited pronounced defects in reversal frequency during the local search (Figure 1A). However, the number of mutants that showed defects in reversal frequency during global search was fewer (Figure 1B). This observation suggests that neuropeptide signalling plays a critical role in regulating the frequency of reversals, particularly during the initial stages of foraging when the organism is actively exploring its immediate environment.

### FLP-15 regulates reversal frequency during foraging

Amongst all the neuropeptide mutants screened, the *flp-15* mutants that show a loss of all isoforms of *flp-15* exhibited an intriguing phenotype (mutant illustrated in Figure S2A). These mutants showed a significant decrease in the frequency of reversals during the local search in comparison to WT animals (Figure 2A). Interestingly, unlike WT *C. elegans*, the *flp-15* mutants failed to reduce their reversal frequency upon transitioning from the local to the global search (Figure 2A). In order to test if the *flp-15* phenotypes were specific to local and global search conditions or if they prevailed in well-fed conditions, we went on to test for the reversal frequency of *flp-15* mutants in the presence of food. Our findings indicated that *flp-15* mutants did not exhibit a significant change in reversal frequency when on food in comparison to WT control animals (Figure 2B). To analyse any possible defect in *flp-15* locomotory speed, we analysed the speed of *C. elegans* during local and global search and found that the *flp-15* mutants moved at speeds comparable to WT during local search but showed slower speeds during global search in comparison to WT control animals (Figure S2B). To further establish the specific role of FLP-15 in regulating reversal behaviour, we performed rescue experiments by supplementing *flp-15* mutants with FLP-15 cDNA plasmid construct (P*flp-15*::*flp-15* cDNA::T2A::GFP) under the control of its endogenous promoter. We were able to completely rescue the *flp-15* mutant phenotypes, restoring the normal pattern of reversal frequency in both local to global search (Figure 2C and D). These results suggest a potential role for FLP-15 in modulating reversals during foraging. We next wanted to study the expression pattern and site of action of FLP-15.

**Figure 2:**
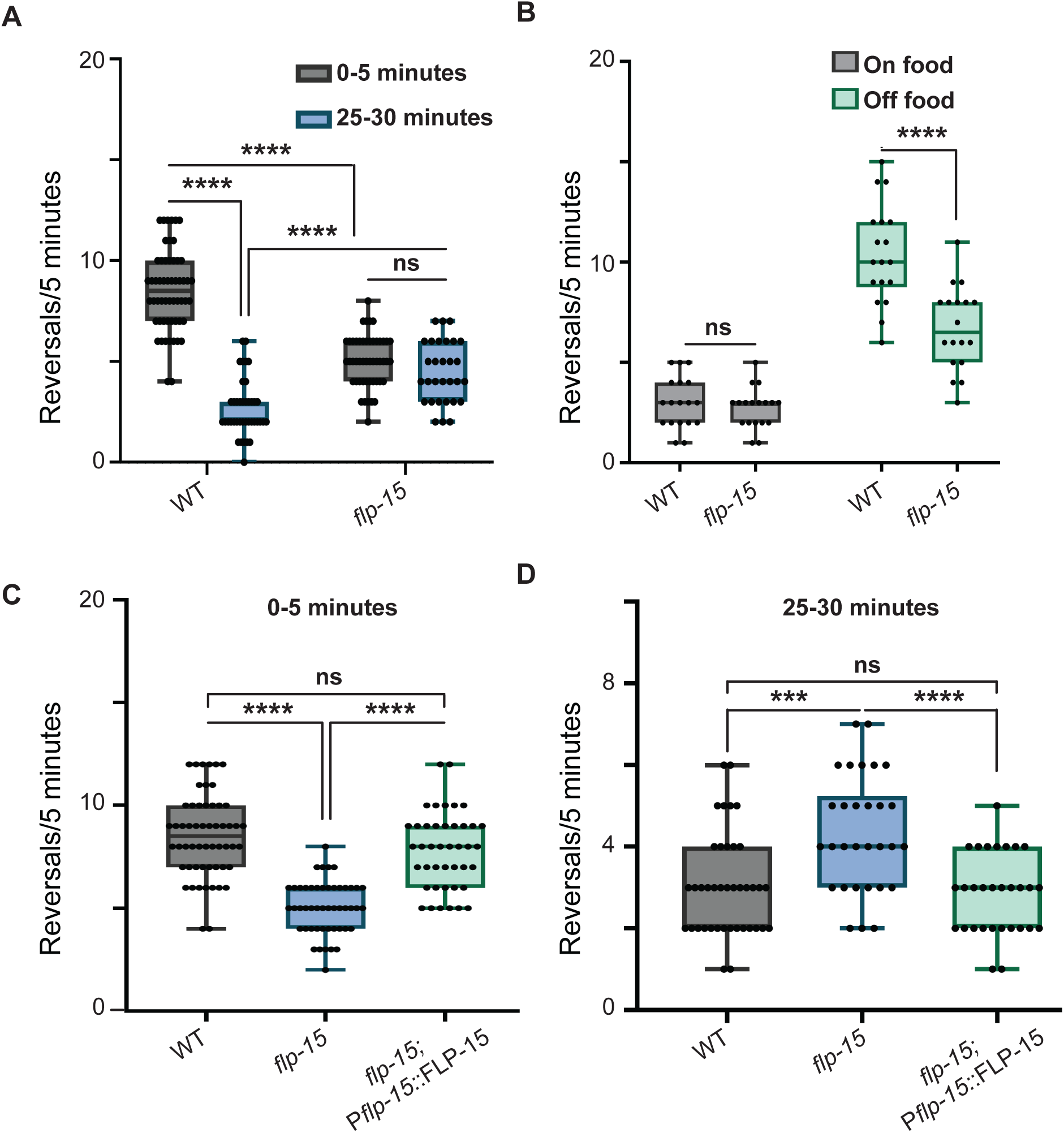
FLP-15 regulates reversal frequency during local and global search. (A) Quantification of reversals in WT and *flp-15* mutants indicating a loss of temporal decay in reversal frequency in *flp-15* mutants during the transition from local to global search. (B) Quantification of reversals in WT and *flp-15* mutants on “on food” conditions indicates no change in reversal frequency in *flp-15* mutants. (C-D) Quantification of reversal frequency with rescue of the phenotype using endogenous promoter of FLP-15 neuropeptide gene. Each dot on the plots corresponds to the number of spontaneous reversals observed per 5 minutes from a single animal. Statistical significance was determined using One-way ANOVA with Tukey test for multiple comparisons, “ns” indicates not significant, “***’’ or “****” indicate p-values less than 0.001 or 0.0001, respectively.

### FLP-15 functions through the I2 pharyngeal neurons

Given that FLP-15 has not previously been implicated in the foraging circuit, we sought to characterize its expression pattern. We generated a transgenic *C. elegans* line carrying a P*flp-15*::GFP reporter construct and observed GFP expression in a single head neuron and a couple of tail neurons (Figures 3A and S3). The head neuron appeared to be the I2 pharyngeal neuron when compared to the anatomical structure available on Wormatlas. To validate this finding, we generated another transgenic line containing both P*flp-15*::GFP and an I2 neuron specific marker, P*gur-3*::mCherry. Co-localization analysis revealed that the two fluorophores indeed co-localized specifically within the I2 neuron (Figure 3A). To test if FLP-15 functions through the I2 neurons, we conducted rescue experiments by supplementing *flp-15* mutant animals with a FLP-15 cDNA plasmid construct (P*gur-3*::*flp-15* cDNA::T2A::GFP) under the control of the I2-specific promoter. Our rescue construct successfully restored the WT pattern of reversal frequency observed during local and global search behaviours in *flp-15* mutant animals (Figure 3B and C).

**Figure 3:**
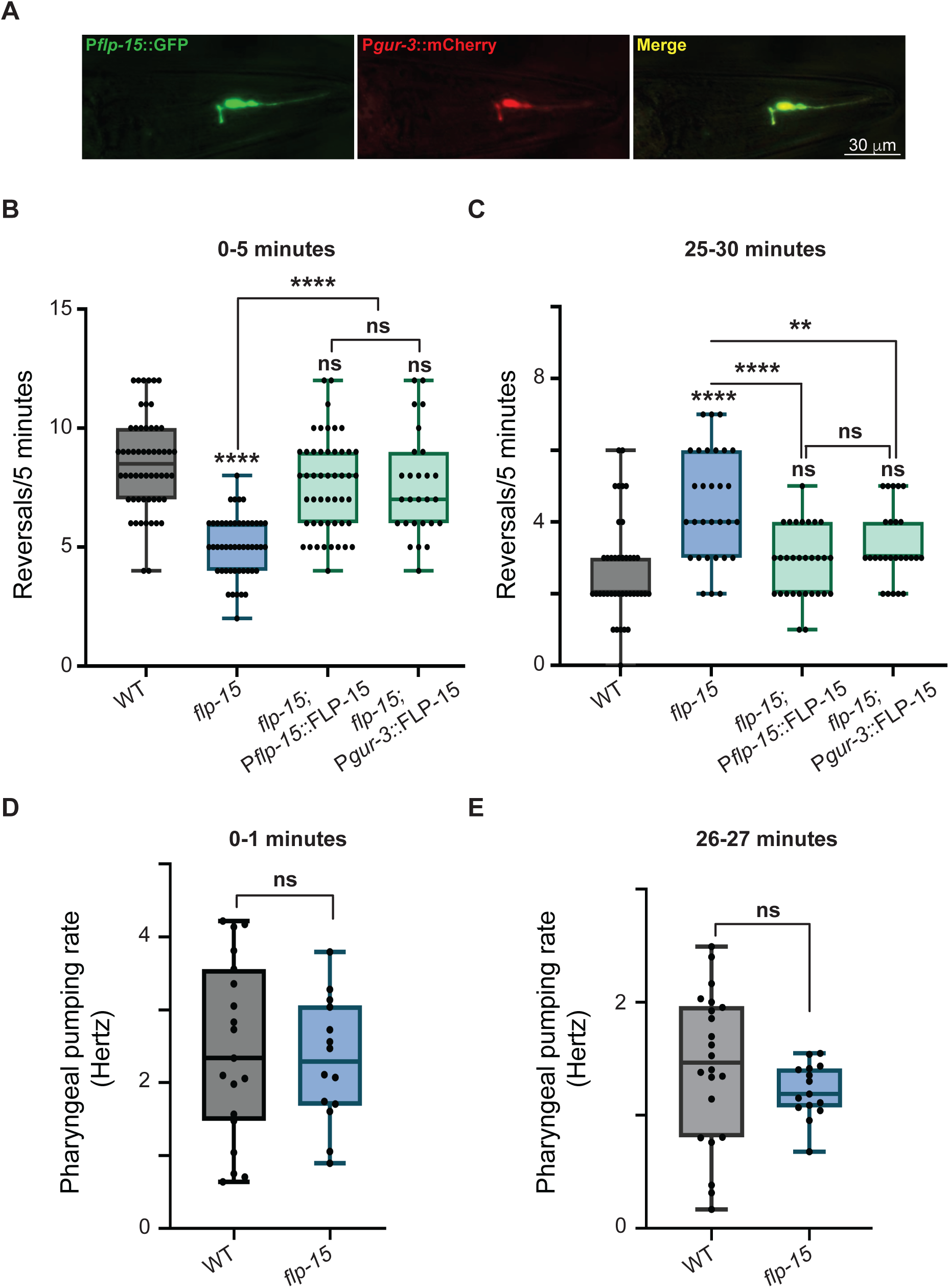
FLP-15 functions through I2 pharyngeal neuron. (A) To elucidate the spatial expression pattern of FLP-15 within the *C. elegans* nervous system, a transgenic line was made by co-injecting a construct containing P*flp-15*::GFP along with another construct containing P*gur-3*::mCherry. Through colocalization analysis, FLP-15 expression was specifically localized to the I2 neuron. (B-C) I2 neuron specific transgenic line carrying the P*gur-3*::FLP-15::GFP construct, introduced into the *flp-15* mutant background successfully restored the decrease in reversal frequency observed during the local and global search. Each dot on the plot corresponds to the number of spontaneous reversals observed per 5 minutes from a single animal. (D-E) Frequency of pharyngeal pumping in WT and *flp-15 C. elegans* shows no differences in pumping rate during local and global search. Statistical analysis was performed using one-way ANOVA with post-hoc Tukey test for multiple comparisons, “ns” indicates not significant, “∗∗” indicates p < 0.01 and “∗∗∗∗” indicates p < 0.0001.

Given the expression of FLP-15 in the I2 pharyngeal neurons, we next went on to test whether the phenotype in *flp-15* mutants is associated with defects in pharyngeal pumping. Our results show that there is no significant change in the rate of pharyngeal pumping between *flp-15* and WT animals (Figure 3D and E). These results provide compelling evidence that the expression of FLP-15 in the I2 neuron is indeed sufficient to rescue the *flp-15* mutant phenotype, indicating that FLP-15 functions from the I2 neuron to regulate reversal frequency during foraging in *C. elegans*. Further, the foraging behaviour in *flp-15* mutants appears to be uncoupled from pharyngeal pumping as *flp-15* mutants that show defects in both local and global foraging behaviours show no defects in pharyngeal pumping at similar time scales (Figure 3D and E).

Having elucidated the site of action of FLP-15, we were interested in finding the receptor through which FLP-15 may be functioning to regulate foraging behaviour in *C. elegans*

### FLP-15 functions through the NPR-3 receptor to regulate reversal frequency

Our next set of experiments aimed to identify the receptor responsible for mediating the effects of the neuropeptide FLP-15 in *C. elegans* foraging. Through a review of the existing literature, we found NPR-3 as the receptor known to exhibit strong one-to-one binding with FLP-15 (Beets *et al*, 2023; Gershkovich *et al*, 2019). However, due to the unavailability of a *npr-3* mutant strain we utilized the CRISPR-Cas9 technology to generate a null mutant of *npr-3* (Figure S4). This mutant has a deletion spanning 2.8 kb, encompassing the region from exon 1 to exon 6 of the *npr-3* gene (illustrated in Figure 4A). Locomotion assays were conducted with these mutants to assess the behavioural phenotype of the *npr-*3 mutant animals. We found significant alterations in the foraging behaviours in *npr-3* mutants when compared to WT control animals. Specifically, *npr-3* mutant *C. elegans* exhibited a notable decrease in the frequency of reversals during local search behaviour. Additionally, they failed to demonstrate the temporal decrease in reversal frequency during the transition from local to global search. Overall *npr-3* mutants showed phenotypes that were indistinguishable from that of *flp-15* mutant *C. elegans* (Figure 4B and C). To confirm that these phenotypic changes were due to the loss of *npr-3* function, we conducted rescue experiments by microinjecting the genomic DNA of the *npr-3* gene under the control of its endogenous promoter into the *npr-3* mutant *C. elegans.* The reversal frequency in these transgenic animals (*npr-3*; P*npr-3*::npr-3::GFP) were similar to that of WT animals, suggesting successful restoration of NPR-3 function (Figure 4B and C). To further elucidate the genetic pathway underlying the observed phenotype, we investigated the potential genetic interaction between *flp-15* and *npr-3*. Given the similarity in locomotion defects between *flp-15* and *npr-3* mutants, we hypothesized that FLP-15 functions through NPR-3 to regulate reversal frequency during foraging. To test this hypothesis, we generated *flp-15; npr-3* double mutants and conducted locomotion assays with these animals. The defects in the frequency of reversals observed in the *flp-15; npr-3* double mutants closely resembled that of *flp-15* and/or *npr-3* single mutant animals (Figure 4B and C), providing strong evidence that FLP-15 acts through NPR-3 in regulating reversal behaviour during foraging in *C. elegans*.

**Figure 4:**
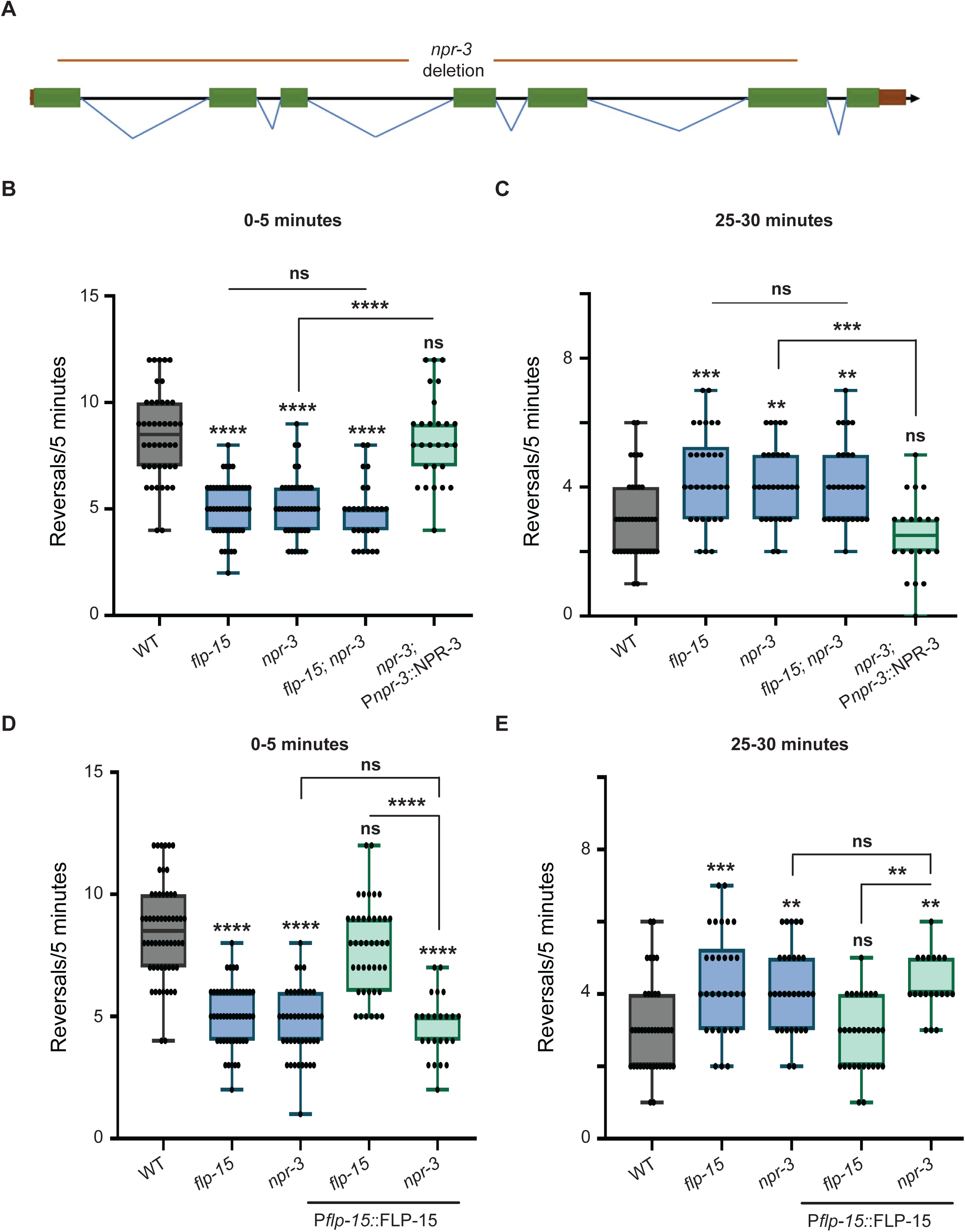
NPR-3 is the receptor for FLP-15. (A) Schematic of the *npr-3* CRISPR mutation used in this study. (B-C) Reversal frequency during local and global search in *flp-15*, *npr-3*, *flp-15; npr-3* and *npr-3* rescue animals. In (B) the reversal frequency shows significant reduction during local search in both *flp-15* and *npr-3* mutants. A similar defect in reversal frequency was found in the *flp-15; npr-3* double mutants. The phenotype in *npr-3* mutants was rescued by supplementing *npr-3* gene under the control of its endogenous promoter. (C) indicates reversal frequency during global search which was significantly increased in *flp-15, npr-3* mutants and the *flp-15; npr-3* double mutants. The phenotype in *npr-3* mutants was rescued by supplementing the *npr-3* gene under the control of its endogenous promoter. (D-E) Overexpression of the neuropeptide FLP-15 in the *npr-3* mutant background in unable to alleviate the defect in reversal frequency in these mutants during local and global search, indicating NPR-3 in required for FLP-15 mediated regulation of reversal frequency. Each dot on the plots corresponds to the number of spontaneous reversals observed per 5 minutes from a single animal. Statistical significance was determined using One-way ANOVA with Tukey test for multiple comparisons, “ns” indicates not significant, “**”, “***’’ or “****” indicate p-values less than 0.01, 0.001 or 0.0001, respectively.

In order to further strengthen the evidence supporting the genetic interaction between *flp-15* and *npr-3*, we overexpressed neuropeptide FLP-15, by micro injecting the plasmid construct containing P*flp-15::*FLP-15::T2A::GFP, in the *npr-3* mutant background and subsequently conducted locomotion assays. Interestingly, our results revealed that *npr-3* mutant animals overexpressing FLP-15 displayed a locomotion phenotype similar to that observed in both *flp-15* and *npr-3* mutants (Figure 4D and E). This finding strongly suggests that NPR-3 serves as the receptor for FLP-15, mediating its regulatory role in controlling the frequency of reversals during foraging. We were next interested in identifying the neurons through which NPR-3 functions.

### NPR-3 is expressed in dopaminergic neurons

We explored the expression pattern and potential neuron specific role of NPR-3 in locomotion, as no prior investigations had delved into these aspects. To this end, we used a plasmid construct containing P*npr-3*::GFP and expressed it in WT *C. elegans*. Our analyses revealed NPR-3 expression in several head neurons and a single tail neuron (Figure 5A). This indicates a potentially widespread distribution of NPR-3 within the nervous system. To further characterize the identity of these neurons, we searched through the *C. elegans* single-cell RNA sequencing (scRNAseq) databases, CeNGEN and VISTA (Hammarlund *et al*, 2018; Liska *et al*, 2023). The data from these databases provided insights suggesting NPR-3 expression in dopaminergic neurons and various other head neurons, as well as in the DVA neuron located in the tail region.

**Figure 5:**
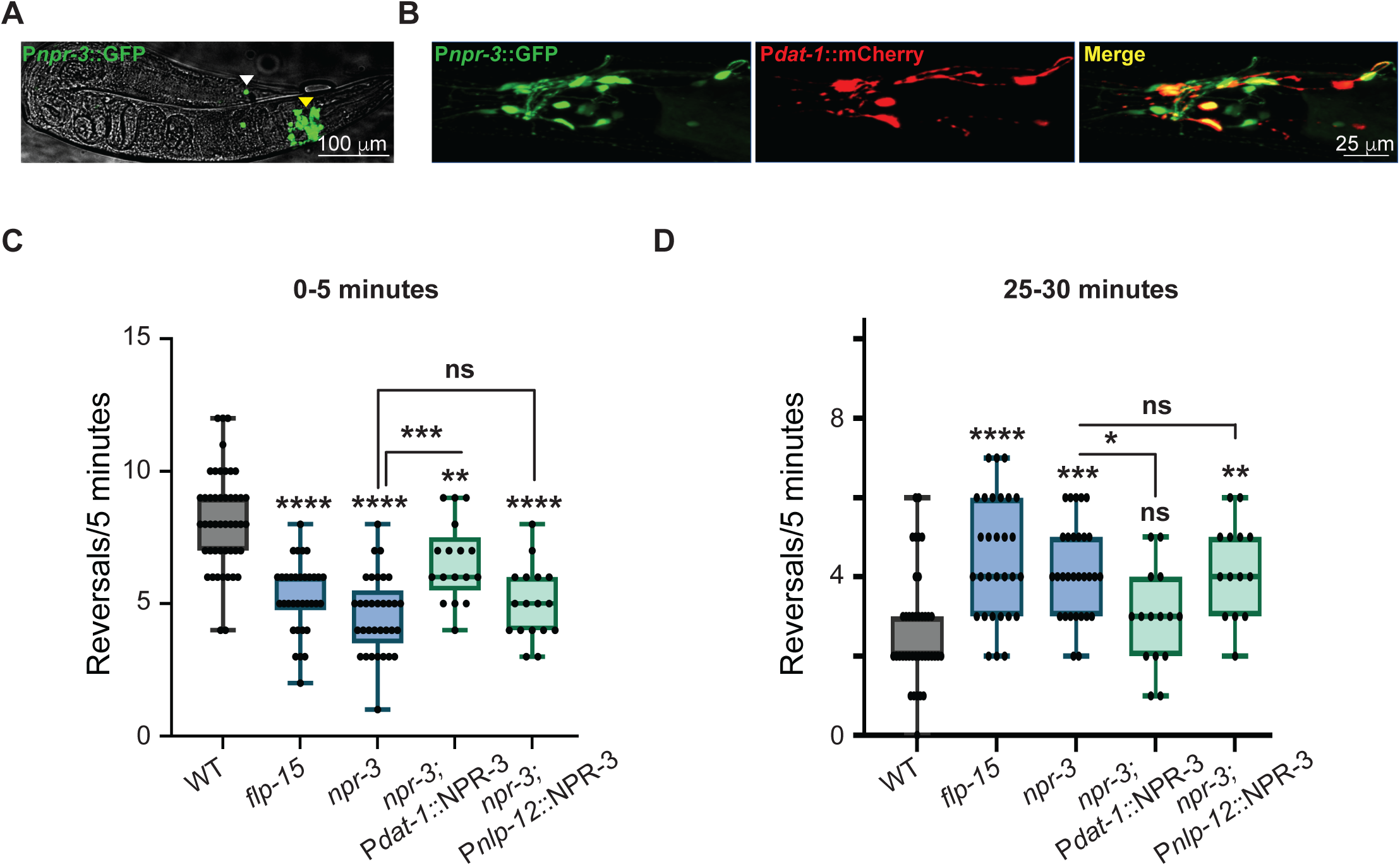
NPR-3 is expressed in head and tail neurons. (A) P*npr-3*::GFP shows expression in multiple head neurons and a tail neuron. (B) Spatial expression pattern of NPR-3 within the *C. elegans* head neurons shows expression in multiple neurons. The expression of NPR-3 is seen to partially colocalize with dopaminergic neurons using P*dat-1*::mCherry for colocalization with P*npr-3*::GFP. (C-D) Quantification of spontaneous reversals in *npr-3* mutants with NPR-3 expressed in dopaminergic and DVA neurons during local and global search. NPR-3 shows rescue when expressed in dopaminergic neurons and not the DVA neuron. Each dot on the plots corresponds to the number of spontaneous reversals observed per 5 minutes from a single animal. Statistical analysis was performed using one-way ANOVA with post-hoc Tukey test for multiple comparisons, “ns” indicates not significant, “*”, “**”, “***’’ or “****” indicate p-values less than 0.05, 0.01, 0.001 or 0.0001, respectively.

Since multiple head neurons expressing NPR-3 included dopaminergic neurons, we sought to confirm the expression of NPR-3 in dopaminergic neurons specifically. We generated a transgenic line by expressing of mCherry under the control of a dopaminergic neuron-specific promoter, *dat-1*. Colocalization analysis with dopaminergic neurons revealed NPR-3 expression in six dopaminergic neurons in the head, comprising two pairs of CEP neurons and one pair of ADE neurons (Figure 5B). The anatomical positioning and structural characteristics of the tail neuron further supported the expression of NPR-3 in the DVA neuron (Figure 5A). We were unable to localise the expression of NPR-3 in the remaining head neurons due to unavailability of neuron specific promoters for those putative neurons. These findings collectively provide a comprehensive understanding of NPR-3 expression patterns within the nervous system of *C. elegans*, particularly highlighting its presence in dopaminergic neurons and the DVA neuron.

To further delve into the neuron-specific role of NPR-3 in regulating reversals, we conducted rescue experiments targeting dopaminergic and DVA neurons in *npr-3* mutant *C. elegans*. By introducing a genomic DNA fragment of NPR-3 under the control of the *dat-1* promoter, we specifically expressed NPR-3 in dopaminergic neurons. In addition to this experiment, we also expressed NPR-3 under the *nlp-12* promoter to express NPR-3 in DVA neurons. Upon quantifying spontaneous reversals during both local and global search behaviours in the rescued *npr-3* mutant animals we found that expressing NPR-3 specifically in dopaminergic neurons largely rescued the mutant phenotype while expressing NPR-3 in the DVA neuron was insufficient to rescue the reversal defects in the mutant animals (Figures 5C and D). These data indicate that dopaminergic neurons play a pivotal role in mediating the effects of NPR-3 regulation of foraging. We next wanted to explore the role of dopamine in the FLP-15/NPR-3 pathway.

### FLP-15 regulates dopamine level during foraging

To investigate the role of dopaminergic signalling in the modulation of foraging behaviours, we performed locomotion assays in *cat-2* mutant animals, which lack tyrosine 3-monooxygenase, an enzyme essential for dopamine biosynthesis. These mutants are known to have a 60–70% reduction in dopamine levels (Sanyal *et al*, 2004). Our results indicate that *cat-2* mutants exhibit a significantly larger number of reversals during both local and global search when compared to WT animals in the corresponding behavioural states (Figures 6A and B). However, despite this elevated reversal activity, *cat-2* mutants still display a reduction in reversal frequency during the transition from local to global search (Figure S6). These data suggest that a decrease in dopamine levels may be required to regulate the frequency of reversals during local and global search behaviours. To explore whether FLP-15 functions in this dopaminergic pathway, we analysed reversal behaviours in *cat-2; flp-15* double mutants. Interestingly *cat-2* mutants appear to suppress the *flp-15* phenotype during local search, while during global search the reversal frequencies were comparable to *flp-15* mutants (Figure 6A and B). These results indicate that FLP-15 may act through the dopaminergic pathway.

**Figure 6:**
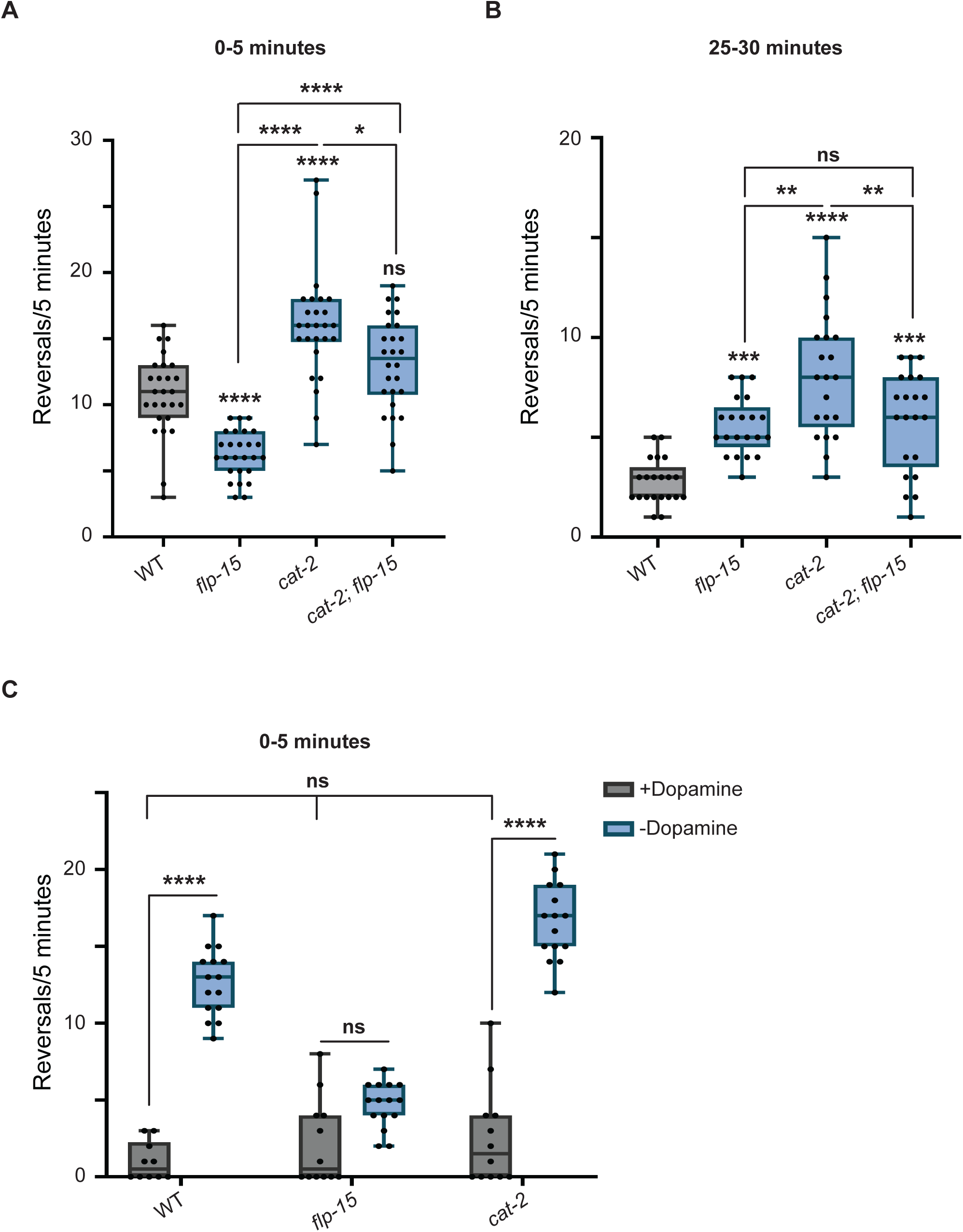
FLP-15 possibly regulates dopamine level during exploration. (A) Quantification of reversals during local search in dopamine synthesis mutants indicates that loss of dopamine leads to increased reversal frequency in *cat-2* mutants. (B) Quantification of reversals during global search indicates that while *cat-2* mutants are able to show a temporal decline in the frequency of reversals, *flp-15* mutants fail to do so. (C) A plot compares reversal frequency between WT, *flp-15*, and *cat-2* mutants with and without dopamine supplementation. Exogenous dopamine supplementation significantly reduced reversal frequency in WT and *cat-2* mutants, indicating enhanced dwelling behaviours. However, the difference in the reversal frequency in *flp-15* mutants was not significant. Each dot on the plots corresponds to the number of spontaneous reversals observed per 5 minutes from a single animal. Statistical analysis was performed using one-way ANOVA with post-hoc Tukey test for multiple comparisons, “ns” indicates not significant, “*”, “**”, “***’’ or “****” indicate p-values less than 0.05, 0.01, 0.001 or 0.0001, respectively.

Previous studies have shown that dopamine promotes dwelling behaviour in *C. elegans* in the presence of food, characterized by decreased speed and a reduced area of exploration (Oranth *et al*., 2018; Sawin *et al*, 2000). In contrast, during extended food deprivation, the absence of dopamine alleviates dwelling behaviour, promoting roaming and increased exploratory activity (Oranth *et al*., 2018). This behavioural transition aligns with the foraging model, which proposes that a gradual decline in reorientation events, such as reversals, is essential for the shift from local to global search strategies (Calhoun *et al*, 2014; Lopez-Cruz *et al*., 2019). We observed that *cat-2* mutant *C. elegans* exhibited increased reversal frequency and enhanced exploratory behaviour, consistent with the roaming state. On the other hand, *flp-15* mutants displayed a reduced reversal frequency and slow exploration, resembling a dwelling-like state in the absence of food. To further investigate the role of dopamine and FLP-15 in this behaviour, we supplemented WT, *cat-2* and *flp-15* mutants with exogenous dopamine and recorded their behaviour. Our results demonstrate that dopamine supplementation led to a reduction in reversal frequency across all genotypes, supporting the hypothesis that elevated dopamine signalling induces slower exploratory behaviour characterised by reduced reversal frequency (Figure 6C). While the decrease in reversals was statistically significant in WT and *cat-2* mutants, *flp-15* mutants did not show a significant change in reversal frequency upon dopamine treatment (Figure 6C). These findings suggest that FLP-15 may be required for effective dopaminergic modulation of food-seeking behaviour, potentially by influencing dopamine signal transmission.

## Discussion

The nervous system orchestrates the intricate coordination between internal state and external environment of an organism, ultimately driving specific behaviors (reviewed in (Chen *et al*, 2021)). Central to this process is the sequential communication among a defined subset of neurons. In this study, we delve into the regulatory role of a neuropeptide, FLP-15, released from the I2 pharyngeal neuron to control various facets of locomotion during foraging. Several reports have suggested the role of neuropeptides in regulating the reversal behavior during exploration for food ((Bhardwaj *et al*., 2018; Cohen *et al*, 2009; Oranth *et al*., 2018) and reviewed in (Bhat *et al*., 2021; Watteyne *et al*., 2024)). However, FLP-15 has not been studied in any aspect of locomotion previously. Our findings underscore the indispensable nature of the FLP-15 neuropeptide in modulating reversal frequency during foraging behavior. Notably, *flp-15* mutants exhibited a significant reduction in reversal frequency during local search compared to wild-type (WT) *C. elegans*, indicative of the pivotal role of FLP-15 in this locomotory aspect.

Interestingly, our results also revealed a notable defect in reversal frequency dynamics between *flp-15* mutants and WT animals during the transition from local to global search. While WT *C. elegans* exhibited a decrease in reversal frequency during this transition, indicative of efficient exploration strategy adaptation, *flp-15* mutants failed to demonstrate this decrease. This highlights the critical role of FLP-15 in orchestrating the temporal dynamics of reversal frequency, a key determinant in shaping the exploration strategy during foraging in *C. elegans* (Model illustrated in Figure 7). It’s worth noting that the frequency of reversals holds paramount importance not only in foraging but also in other vital behaviors such as chemotaxis, mating and pathogen avoidance, where modulation of locomotory strategies is imperative for navigating toward food sources or away from potential threats ((Pierce-Shimomura *et al*, 1999; Sawin *et al*., 2000; Zhao *et al*, 2003) and reviewed in (Bargmann & Mori, 1997)).

**Figure 7:**
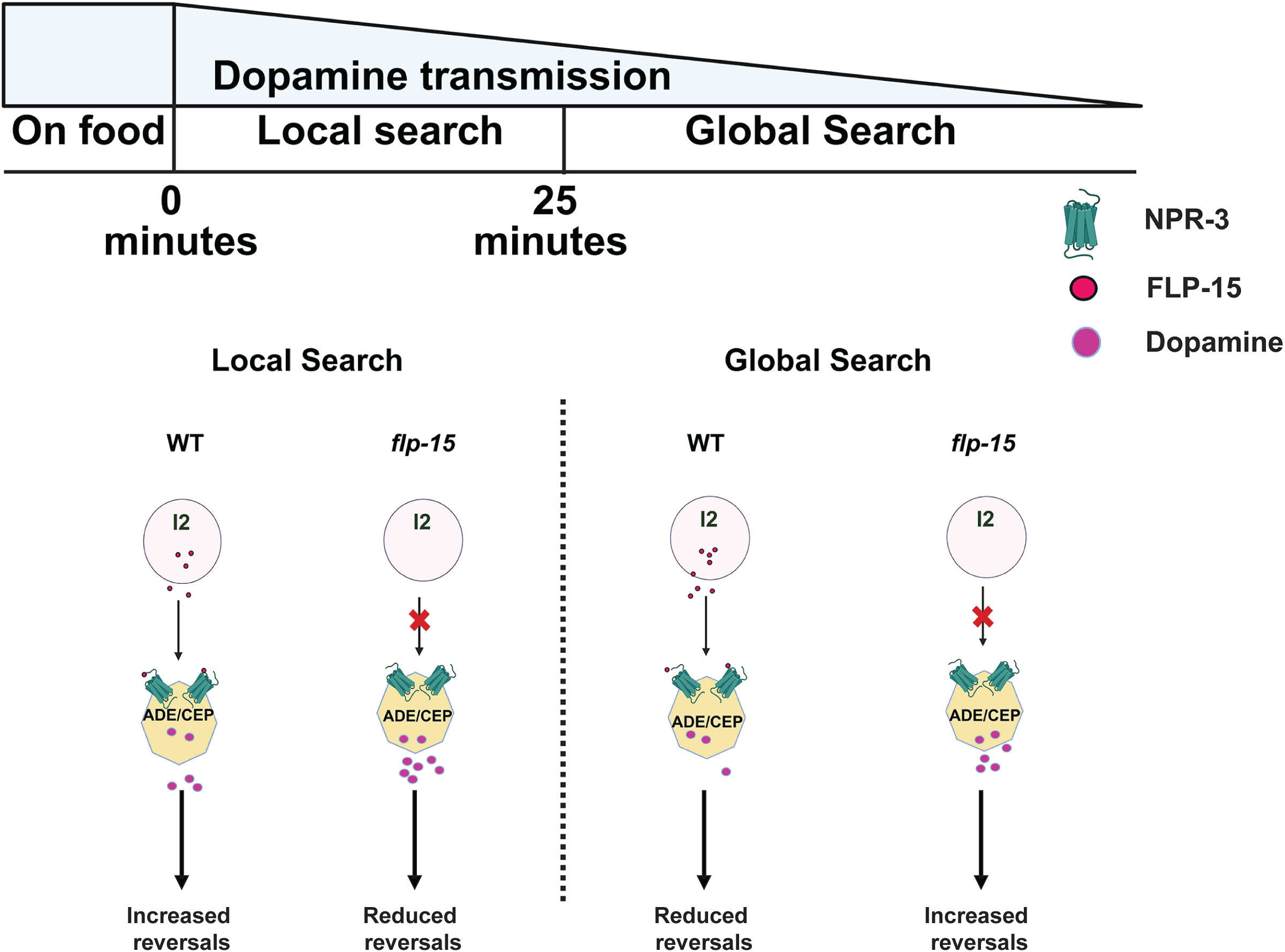
Model for FLP-15 function. The illustration shows the proposed mechanism of FLP-15/NPR-3 mediated regulation of locomotion during foraging. We propose that gradual decline in dopamine signalling enables the *C. elegans* to transition from local search to global search behaviours. FLP-15 released from the I2 pharyngeal neuron, modulates the dopaminergic transmission to regulate the frequency of reversals and enables the efficient foraging during food seeking behaviours.

Further results from this study suggest that FLP-15 functions through the I2 pharyngeal neuron to regulate the frequency of reversals during both local and global search behaviors. Interestingly, we did not find any significant defect in the pharyngeal pumping rate in *flp-15* mutants. Previous studies have indicated that the rate of pharyngeal pumping in *C. elegans* correlates with recent food encounters, with pumping rates decreasing over time in response to extended periods of starvation or off-food conditions (Bonnard *et al*., 2022; Scholz *et al*, 2016). The correlation between pharyngeal activity and foraging behavior underscores the intricate interplay between internal metabolic cues and locomotor responses, highlighting the significance of pharyngeal function in modulating the dynamics of foraging strategies in *C. elegans*. However, the data from this study indicates that FLP-15 does not have any role in regulating the rate of pharyngeal pumping. Our findings also establish NPR-3 as the key receptor responsible for mediating the effects of FLP-15 on reversal frequency during foraging behavior. NPR-3 has been characterized as an inhibitory G_i_ protein (Gershkovich *et al*., 2019; Kubiak *et al*, 2003). This indicates that FLP-15 binding to NPR-3 may lead to inhibition of activity of the neuron expressing NPR-3. Previous studies have extensively delved into the role of dopamine in modulating the locomotion in the context of food (Hills *et al*, 2004; Oranth *et al*., 2018). These studies are consistent in establishing that dopamine is required to maintain dwelling behavior on food and subsequently in off food condition the dopamine signaling is reduced to enable roaming behaviours. Oranth *et al* have shown that photostimulation of ADE and CEP neurons in off food condition leads to a slowing down response in *C. elegans* and mimics basal slowing behavior (Oranth *et al*., 2018). Consistent with these data, our study indicates that loss of FLP-15/NPR-3 signaling in mutants may result in the activation of ADE and/or CEP neurons and as a result show a reduced frequency of reversals during local search when compared to their WT counterparts where the reduced dopaminergic signaling ensures increased frequency of reversals. Moreover, our observation that NPR-3 primarily functions through dopaminergic neurons, rather than DVA neurons, in regulating reversal frequency provides important insights into the neural circuitry governing foraging behavior in *C. elegans*. This suggests a potential interaction between NPR-3 and dopamine signaling pathways, implicating a coordinated regulatory mechanism in modulating reversal frequency during both local and global search behaviors. The expression pattern of NPR-3 and rescue in specific subsets of dopaminergic neurons offers valuable clues regarding its potential role in modulating locomotion behavior mediated by dopamine. Dopaminergic signaling has been extensively implicated in motor control and behavioral regulation across various species ((Omura *et al*, 2012; Pandey *et al*., 2021) and reviewed in (Beninger, 1983; Ryczko & Dubuc, 2017; Sharples *et al*, 2014)), suggesting a plausible coordinating role of NPR-3 and dopamine in regulating reversal frequency during foraging. Furthermore, the expression of NPR-3 in the DVA neuron, which is known for its role in coordinating locomotion and sensory processing in the tail region (Bhattacharya *et al*, 2014; Hu *et al*, 2011; Pandey *et al*., 2021; Ramachandran *et al*., 2021), suggests a broader functional significance of NPR-3 beyond the head neurons. This implies that NPR-3 may mediate a multifaceted role in modulating locomotion and sensory integration throughout the nervous system of *C. elegans*. These observations open new avenues for exploring downstream effectors and signaling pathways involved in NPR-3-mediated regulation of reversal behavior and neural circuitry.

Our findings from the exogenous dopamine supplementation assay further support the role of dopaminergic modulation in regulating exploratory behavior in *C. elegans*. Irrespective of the genotype, *C. elegans* supplemented with exogenous dopamine predominantly exhibited dwelling behavior, in contrast to their counterparts with no dopamine supplementation, which displayed roaming behavior. Notably, *cat-2* mutant *C. elegans*, which are deficient in dopamine synthesis (Sanyal *et al*., 2004), showed a significant decrease in the number of reversals upon dopamine supplementation. In contrast, although *flp-15* mutant animals appeared to exhibit a reduced reversal frequency with dopamine supplementation, this reduction was not significantly different from the no dopamine supplemented *flp-15* mutants. This suggests that *flp-15* mutants may already have elevated dopamine signaling, resulting in a constitutive dwelling-like behavioral state even in the absence of exogenous dopamine. This implies that FLP-15 mediated modulation of locomotion behavior may involve complex interactions with dopaminergic signaling pathways, highlighting the need for further investigations to elucidate the underlying mechanisms. Growing evidence supports the role of dopamine in regulating reward-based food intake and appetite control (reviewed in (Baik, 2013, 2021)). A deeper understanding of the underlying molecular mechanisms may facilitate the development of targeted therapies for metabolic disorders linked to dysregulated feeding behavior and appetite control (reviewed in (Volkow *et al*, 2011)).

## Acknowledgements

The authors are grateful to Bill Schafer for a strain. Some strains were provided by CGC, which is funded by NIH Office of Research Infrastructure Programs (P40 OD010440). All illustrations were created in BioRender. We thank Faraz Abbas, Imra Aftab and Velu Saravanan for help with experiments. We also thank Palagiri Suresh for routine help and the members of Kavita Babu’s lab for suggestions and critique on the manuscript.

## Author Contributions

USB: Design and execution of experiments, data analyses, manuscript writing/editing, SS: Outcrossing and behavioural screen, SH: Molecular biology and behavioural experiments and manuscript editing, JL: strain crossing and genotyping, YX: Genotyping and behavioural experiments, NT: Molecular biology experiments, AB: Conceptualizing the screen, MS: data analysis, and KB: Supervision, funding acquisition and manuscript writing/editing.

