## Supplemental information for "FLP-15 functions through the GPCR NPR-3 to regulate local and global search behaviours in *Caenorhabditis elegans*"

**Supplementary Information:**

**Table S1 (List of strains)**

| Strain Name | Mutant | CGC Strain | Outcross Status | Figure No. |
|---|---|---|---|---|
| BAB6013 | *flp-33 (gk1037)* | VC2422 | Outcrossed 3X | 1 |
| BAB6014 | *flp-8* (pk360) | PT501 | Outcrossed 3X | 1 |
| BAB6015 | *flp-15* (gk1186) | VC2504 | Outcrossed 3X | 1-6 |
| BAB6016 | *flp-17* (ok3587) | RB2575 | Outcrossed 3X | 1 |
| BAB6017 | *nlp-49* (gk546875) | From William Schafer Lab | | 1 |
| BAB6018 | *flp-2* (ok3351) | VC2591 | Outcrossed 3X | 1 |
| BAB6019 | *flp-6* (ok3056) | VC2324 | Outcrossed 3X | 1 |
| BAB6020 | *flp-20* (pk1596) | PT505 | Outcrossed 3X | 1 |
| BAB6021 | *flp-9* (ok2730) | RB2067 | Outcrossed 3X | 1 |
| BAB6022 | *ins-3* (ok2488) | RB1915 | Outcrossed 3X | 1 |
| BAB6023 | *pdf-1* (tm1996) | LSC27 | Outcrossed 3X | 1 |
| BAB6024 | *flp-5* (gk3123) | Obtained from VC3280 | Outcrossed 3X | 1 |
| BAB6025 | *flp-3* (ok3265) | VC2497 | Outcrossed 3X | 1 |
| BAB6026 | *nlp-11* (syb530) | PHX530 | Outcrossed 3X | 1 |
| BAB6027 | *nlp-8* (ok1799) | VC1309 | Outcrossed 3X | 1 |
| BAB6028 | *ins-4* (ok3534) | RB2544 | Outcrossed 3X | 1 |
| BAB6029 | *nlp-3* (ok2688) | RB2030 | Outcrossed 3X | 1 |
| BAB6030 | *nlp-14* (ok1517) | VC1108 | Outcrossed 3X | 1 |
| BAB6031 | *nlp-15* (ok1512) | VC1063 | Outcrossed 3X | 1 |
| BAB6032 | *flp-12* (ok2409) | RB1863 | Outcrossed 3X | 1 |
| BAB6033 | *nlp-18* (ok1557) | RB1372 | Outcrossed 3X | 1 |
| BAB6034 | *flp-7* (ok2625) | RB1990 | Outcrossed 3X | 1 |
| BAB6001 | *npr-3* (ind6001) | This study | Outcrossed 3X | 4,5 |
| BAB6037 | *flp-15; npr-3* | This study | | 4 |
| BAB6038 | *cat-2* (e1112) | CB1112 | Outcrossed 3X | 6 |
| BAB6039 | *flp-15; cat-2* | This study | | 6 |

| | | | | |
|---|---|---|---|---|
| INF512 | *flp-15*(gk1186) III; gnaIs1 IV | From Monika Scholz lab | | 3 |
| BAB6002 | IndEx6002 [*Pflp-15*::FLP-15cDNA :: T2A :: GFP ; *flp-15*] | This study | | 2, 3 |
| BAB6003 | IndEx6003 [*Pflp-15*::GFP] | This study | | 3 |
| BAB6004 | IndEx6004 [*Pflp-15*::GFP; *Pgur-3*::mCherry] | This study | | 3 |
| BAB6005 | IndEx6005 [P*npr-3*::NPR-3gDNA::GFP; *npr-3*] | This study | | 4 |
| BAB6006 | IndEx6006 [*Pnpr-3*::GFP] | This study | | 5 |
| BAB6007 | IndEx6007 [*Pnpr-3*::GFP; *Pdat-1*::mCherry] | This study | | 5 |
| BAB6008 | IndEx6008 [*Pnpr-3::GFP; Pnlp-12*::mCherry] | This study | | 5 |
| BAB6009 | IndEx6009 [*Pflp-15*::FLP-15cDNA::T2A::GFP; *npr-3*] *(*FLP-15 O/E in *npr-3* mutants) | This study | | 4 |
| BAB6010 | IndEx6010 [*Pdat-1*::NPR-3gDNA::mCherry; *npr-3*] | This study | | 5 |
| BAB6011 | IndEx6006 [*Pnlp-12*::NPR-3gDNA::mCherry; *npr-3*] | This study | | 5 |
| BAB6012 | IndEx6006 [*Pgur-3*::FLP-15cDNA :: T2A :: GFP ; *flp-15*] | This study | | 3 |

**Table S2 (List of primers)**

| Genotyping and cloning primers | | | |
|---|---|---|---|
| **Gene** | **Sequence (5'-3')** | **Primer Name** | **Comments** |
| *flp-21* | TCTGATGCGTTTACAGTCGG | USB01 | External Forward |
| | GATAACAGAAATTTGTGCAGAGATCG | USB02 | External Reverse |
| | GCAACCGTTTATCCAAATCGGAGAG | USB03 | Internal Reverse |
| *pdf-1* | AATGTATTGACACTGCTGACTGC | USB04 | External Forward |

| | | | |
|---|---|---|---|
| | CCTTGTGATTTTCCAGGTTATACAAACT | USB05 | External Reverse |
| | CCTATTCCTGAGGTCGCAGTTGAT | USB06 | Internal Reverse |
| *flp-6* | TGGCTTTTGACTCAGGTTAGGATC | USB07 | External Forward |
| | GCACTTTTTCTCTTTTCCATTGGCA | USB08 | External Reverse |
| | TCCGAAACGCATATACGCTGATTTAC | USB09 | Internal Reverse |
| *flp-12* | GATCTTGAAAATAGTGTGCCTCGTTTTC | USB10 | External Forward |
| | GTCCTCATCGAGCTCGATCTTTTTGA | USB11 | External Reverse |
| | CTATCTGCTCTTTTCCTCGTCCTTCT | USB12 | Internal Reverse |
| *ins-1* | TAGTTATATGGCGTGCGCTCATTTC | USB13 | External Forward |
| | CAAACTTTAAAAGTGACAACATCCATTTGGT | USB14 | External Reverse |
| | GCTAAAAGGGTTGTTGTGAGACGTGAT | USB15 | Internal Reverse |
| *flp-2* | GATCTTGAAAATAGTGTGCCTCGTTTTC | USB16 | External Forward |
| | GTCCTCATCGAGCTCGATCTTTTTGA | USB17 | External Reverse |
| | CTATCTGCTCTTTTCCTCGTCCTTCT | USB18 | Internal Reverse |
| *flp-1* | GTCCTCGTCAAGCACGTC | USB19 | External Forward |
| | CTCGATGCTCTCCACCATGG | USB20 | External Reverse |
| | AACCGGCTTTCTTTCCCAGCG | USB21 | Internal Reverse |
| *nlp-15* | GGCAAAGACGGCAAAGTTTAGTTAACAA | USB24 | External Forward |

| | | | |
|---|---|---|---|
| | GTCCAGCCAGTGAATCGAAAG | USB25 | External Reverse |
| | CTGGACTTTGCTCAAAAACCGTTAG | USB26 | Internal Reverse |
| *flp-10* | AACAATTCAATTTGCCACGTCATCAA | USB27 | External Forward |
| | CAAACAAAAAAGCATTGGGATGTG | USB28 | External Reverse |
| | TGACGCTTCTCCGATGCGACTT | USB29 | Internal Reverse |
| *ins-3* | GCAGATTCATCAGAAGACCTTTCGCAT | USB30 | External Forward |
| | GACCTAATTCTGTACTCTGCTGTGGTT | USB31 | External Reverse |
| | GCTGCAAGTCTTATGCGTAACTGGAT | USB32 | Internal Reverse |
| *nlp-1* | GCTCCACCCTTTGTTTACATATTC | USB39 | External Forward |
| | ATTCAGAAGCGGAAAGAGCATG | USB40 | External Reverse |
| | CAATTGTGTCCTCCCCCTAAAG | USB41 | Internal Reverse |
| *ins-22* | CTTCAGGATCACGTGAAAAGTCTG | USB46 | External Forward |
| | ACGACTTCCAATTTTGCGAGTATG | USB47 | External Reverse |
| | TCGGTGATGCTTGGATACTCTG | USB48 | Internal Reverse |
| *nlp-3* | GAATCGAAAGACCCTGGAAAACTTG | USB49 | External Forward |
| | GCTAGCTTCATCATCAACTGACTG | USB50 | External Reverse |
| | CCACGGATCAGTAAACCGGATAC | USB51 | Internal Reverse |
| *nlp-18* | CTCCCCCAACAATCCATGATTTCC | USB52 | External Forward |

| | | | |
|---|---|---|---|
| | GTGAAGTGGAATCGGATGATCGAC | USB53 | External Reverse |
| | CTCAAACCTCCAATTTCCGAGTTC | USB54 | Internal Reverse |
| *ins-4* | TGAAACCCCACCGAAAGAATTCTG | USB 58 | External Forward |
| | GAGACGGCAAAAAATGGCAAAAAC | USB 59 | External Reverse |
| | TTGATAAAAAAGACAACCGTCCCTC | USB 60 | Internal Reverse |
| *nlp-49* | CAAGTCGCTAACATTCACATCATGT | USB61 | External Forward |
| | TGGAGCTTCAATTTGTTGTGGAG | USB62 | External Reverse |
| | CTTCCGGGAGAAAGTAAACTGACA | USB63 | Internal Reverse |
| *flp-33* | CAATTTCCTCCCGCAAACAATAG | USB66 | External Forward |
| | TAGTATAAAAGCTGTGCAGCCAG | USB67 | External Reverse |
| | AGATTTAGGCTTAGGCTTAGGCTC | USB68 | Internal Reverse |
| *flp-15* | CAGTGAGTCCGAAATTTTAAACCTGA | USB69 | External Forward |
| | AGCATTCTAAAATTGCAGATTACGAC | USB70 | External Reverse |
| | ACTGTGCTGCATGTGTTTCTATG | USB71 | Internal Reverse |
| *flp-17* | ATGTGCCGACTGAAAGAAGAG | USB72 | External Forward |
| | TGGGAATTTTAACAGGGGGTTG | USB73 | External Reverse |
| | TTCATTTTTAGGGGTCTCACAGTC | USB74 | Internal Reverse |
| *flp-9* | CTCTTGTCCATACGCATTTCCTC | USB75 | External Forward |

| | | | |
|---|---|---|---|
| | TGATGACATCATTCAAAGTCACGAG | USB76 | External Reverse |
| | GCACATTTTCGGAATTGGGGAG | USB77 | Internal Reverse |
| *flp-16* | GAATTCAGGGCGATCCAGTGTC | USB99 | External Forward |
| | CATATCTCATCGGTGTCGATTTAAGAG | USB100 | External Reverse |
| | TCGATGACTTTTTCCTTCCATTTTTG | USB101 | Internal Reverse |
| *flp-7* | CTGTTGCTTGGTCTTGTGCAATC | USB102 | External Forward |
| | CGAGATCAGGCTTTTCTGTAGCTG | USB103 | External Reverse |
| | CAAAACGGACCATCGATGATCTC | USB104 | Internal Reverse |
| *flp-33* | CGAGCGAACCATGAAATTGTGTTC | USB105 | External Forward |
| | CACACAGTATTTACGCATTCCTG | USB106 | External Reverse |
| | GTAATGTGGAAGCGGTCTACTG | USB107 | Internal Reverse |
| *flp-5* | CATCATCTCTAGTTCAGCTCCCAAC | USB108 | External Forward |
| | TGTAATGCAGTGTGTACTCAGTCAC | USB109 | External Reverse |
| | GGAAAGAATGGGTCAAAGTGCATC | USB110 | Internal Reverse |
| *flp-12* | GTTAAGTAGCGATGTGTGAAGTTCTC | USB111 | External Forward |
| | GAATTGGTGTCGTCGTGCTAAG | USB112 | External Reverse |
| | GAGCAACTCGATCATTTCTCAATTG | USB113 | Internal Reverse |
| *npr-3* | GAGAAGGCTCGCTGACATTAATG | USB114 | External Forward |

| | | | |
|---|---|---|---|
| | CGATAGTCGTTTTCAGCAATTGTG | USB115 | External Reverse |
| | GAAACGTGCCGATTCGAATGTTG | USB116 | Internal Reverse |
| *nlp-11* | GTCCTCACCATTCCCCTAGG | USB120 | External Forward |
| | GAATAGGAAGAGGGCGGAGG | USB121 | External Reverse |
| | TCTGATCGACGCTGGAAAGA | USB122 | Internal Reverse |
| *nlp-12* | CATGCTCATCCTCGTATTCGTG | NLP-12_EF | External forward |
| | GTCTGCGTCTCTAACATGTTGAC | NLP-12_ER | External reverse |
| | CAAAGAACTCTGCGTCTCAACTC | NLP-12_IR | Internal reverse |
| P*flp-15* | CACAAGCTTCAATCAATATAGCCGGAAATGCCA | HindIII | UB13 |
| | ACCAGGATCCGTATGTGGGAGACCTTCTTCCA | BamHI | UB14 |
| *flp-15* cDNA | ACCAGGATCCTGGAAGAAGGTCTCCCACATAC | BamHI | UB15 |
| | CACAGGGCCCATTTCATAGAAACACATGCAGCAC | XmaI | UB16 |
| P*gur-3* | ATATGCATGCACATCTCATTGGTTGATTTGATC | SphI | UB25 |
| | CTATGGATCCGAAAACGTGACTGACTAACAC | BamHI | UB26 |
| P*npr-3* | TGTGAAGCTTCTTAATACTGTCCGTCTGCAAG | HindIII | UB33 |
| | CACACCCGGGTCAAAAATTCCAGAAGAGGAGAAC | XmaI | UB34 |
| *npr-3* gDNA | TCATCCCGGGATCCAAAATGGAGGGTGGTC | XmaI | UB35 |
| | ACACGGTACCTCTAACAACCCGGTAGAATCATC | KpnI | UB36 |
| P*dat-1* | ATCTGCATGCTGCCACCGATTTGTACAAATG | SphI | UB43 |
| | CTATGGATCCGGCTAAAAATTGTTGAGATTCGAG | BamHI | UB44 |

### Table S3 (List of plasmids)

| Plasmid | Construct | Backbone |
|---|---|---|
| pBAB6001 | *Pflp-15*::FLP-15cDNA::T2A::GFP | pPD95.75 |
| pBAB6002 | *Pgur-3*::FLP-15cDNA::T2A::GFP | pPD95.75 |
| pBAB6003 | *Pflp-15*::GFP | pPD95.75 |

| | | |
|---|---|---|
| pBAB6004 | *Pgur-3*::mCherry | pPD49.26 |
| pBAB6005 | *Pnpr-3*::NPR-3gDNA::GFP | pPD95.75 |
| pBAB6006 | *Pnpr-3::GFP* | pPD95.75 |
| pBAB6007 | *Pdat-1*::mCherry | pPD49.26 |
| pBAB6008 | *pnlp-12*::mCherry | pPD49.26 |
| pBAB6009 | *Pdat-1*::NPR-3gDNA::mCherry | pPD49.26 |
| pBAB6010 | *Pnlp-12*::NPR-3gDNA::mCherry | pPD49.26 |

### Supplementary figure legends

**Figure S1**: (A) Illustration of the protocol for screening the phenotype. A single young adult *C. elegans* was gently picked using an eyelash with halocarbon oil and placed on a plate without food to remove residual food by allowing it to crawl for 1 minute. The animals was then transferred to an assay plate and 2 minutes were allowed for the worm to adapt, this was followed by recording the locomotion for 5 minutes (local search). After remaining on the same plate for 25 minutes, a second 5-minute recording was taken (global search). (B) Violin plots showing quantification of number of body-bends/reversal during local search in neuropeptide mutants. In this plot, each dot corresponds to the number body-bends/reversal observed per 5 minutes from a single *C. elegans*. Statistical significance was determined using One-way ANOVA with Tukey test for multiple comparisons, "**"or "****" indicate p-values less than 0.01, or 0.0001, respectively. The absence of "*" indicates not significantly different from WT control animals.

**Figure S2:** (A) Illustration of the *flp-15* (*gk1186*) mutation which is a 708 base pair deletion that eliminates the entire third exon, which codes for all three isoforms of the FLP-15 neuropeptide. This likely results in a null mutation that eliminates *flp-15*. (B) Quantification of velocity in WT and *flp-15* mutants. In WT *C. elegans* the velocity increases significantly after the transition from local to global search. In *flp-15* mutants the velocity during local and global search is not significantly different**.** Each dot on the plots corresponds to the number of spontaneous reversals observed per 5 minutes from a single animal. Statistical significance was determined using Two- ANOVA with Tukey test for multiple comparisons, "**"or "****" indicate p-values less than 0.01, or 0.0001, respectively and "ns" stands for non-Significant.

**Figure S3:** FLP-15 gene is expressed in the tail neurons apart from the I2 neuron in the head.

**Figure S4:** DNA gel image shows the *npr-3* deletion mutant generated by CRISPR Cas9. A 2816 bp region spanning exons 1 to 6 of the *npr-3* gene was deleted. The band corresponding to the mutant allele appears lower than the WT band confirming the deletion.

**Figure S6:** Quantification of reversals in *cat-2* mutants during local and global search indicating a temporal decay in reversal frequency in *cat-2* mutants during the transition from local to global search. Each dot on the plots corresponds to the number of spontaneous reversals observed per 5 minutes from a single animal. Statistical significance was determined using One-way ANOVA with Tukey test for multiple comparisons, “****” indicate p-values less than 0.0001.

**Figure S1**

**A**

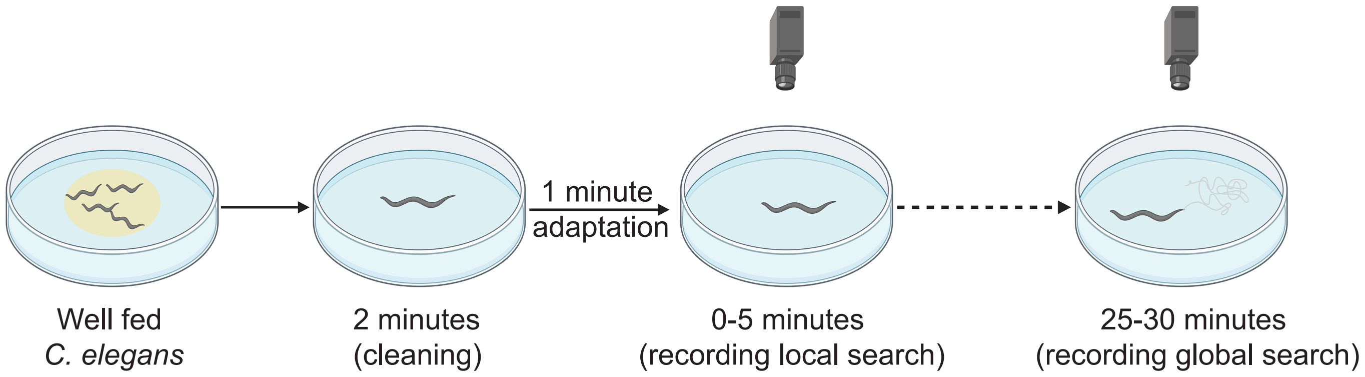

1 minute
adaptation
Well fed
*C. elegans*
2 minutes
(cleaning)
0-5 minutes
(recording local search)
25-30 minutes
(recording global search)


**B**

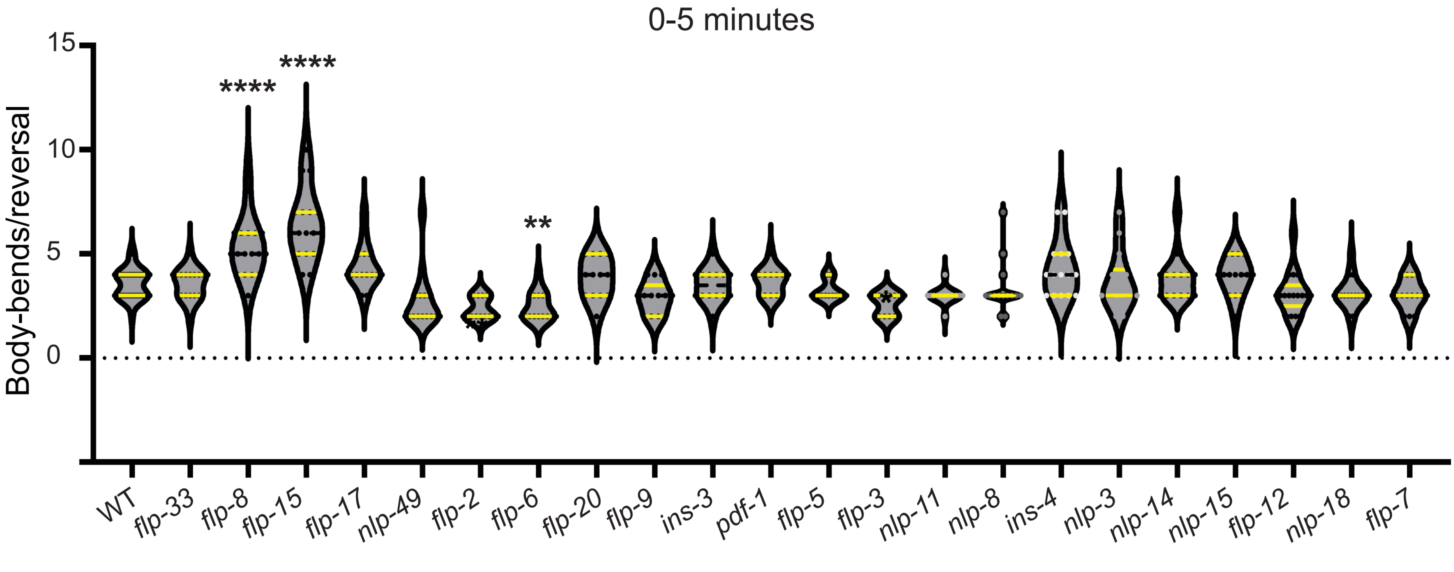

0-5 minutes
Body-bends/reversal
15
10
5
0
****
****
**
WT
flp-33
flp-8
flp-15
flp-17
nlp-49
flp-2
flp-6
flp-20
flp-9
ins-3
pdf-1
flp-5
flp-3
nlp-11
nlp-8
ins-4
nlp-3
nlp-14
nlp-15
flp-12
nlp-18
flp-7

### Figure S2

**A**

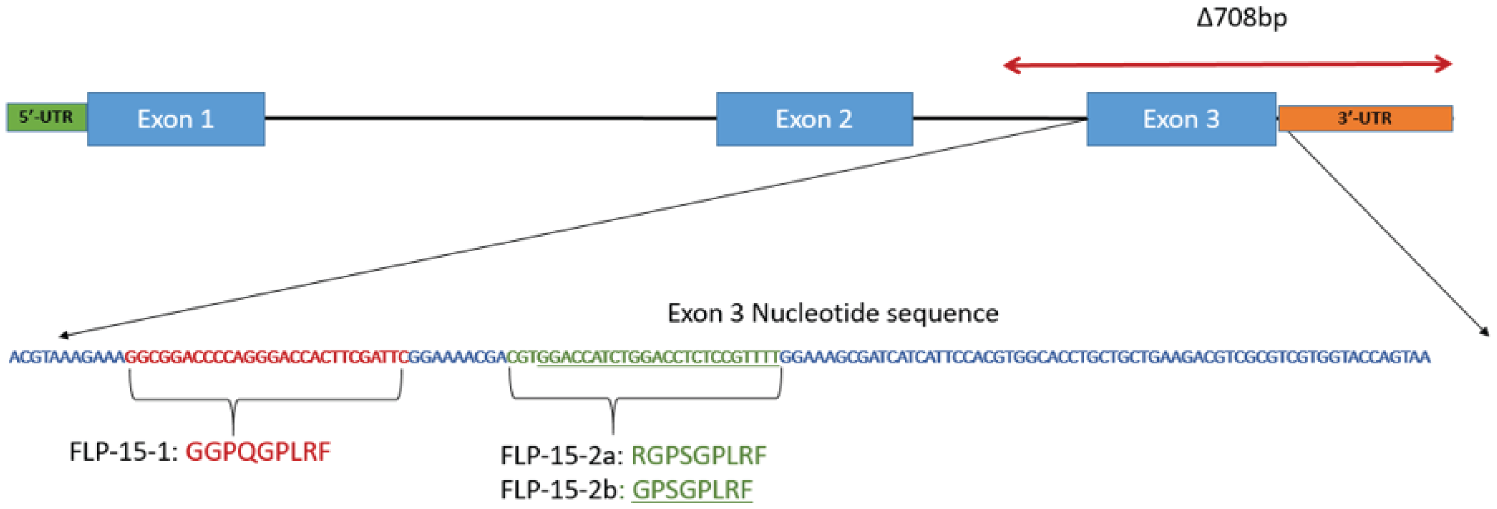
Δ708bp
5'-UTR
Exon 1
Exon 2
Exon 3
3'-UTR
Exon 3 Nucleotide sequence
ACGTAAAGAAAGGCGGACCCCAGGGACCACTTCGATTCGGAAAACGACGTGGACCATCTGGACCTCTCCGTTTTGGAAAGCGATCATCATTCCACGTGGCACCTGCTGCTGAAGACGTCGCGTCGTGGTACCAGTAA
FLP-15-1: GGPQGPLRF
FLP-15-2a: RGPSGPLRF
FLP-15-2b: GPSGPLRF

**B**

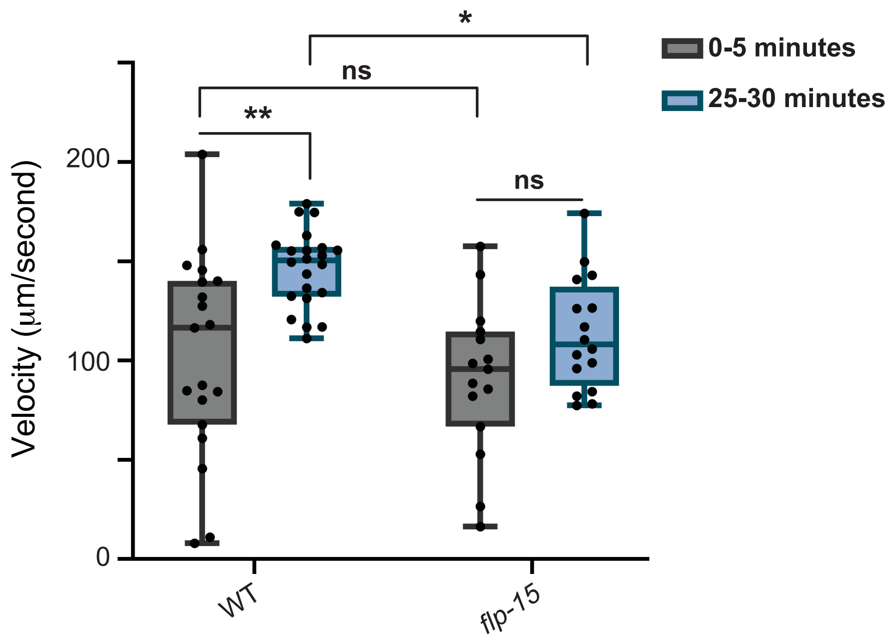
Velocity (μm/second)
0
100
200
**
ns
*
ns
0-5 minutes
25-30 minutes
WT
flp-15

### Figure S3

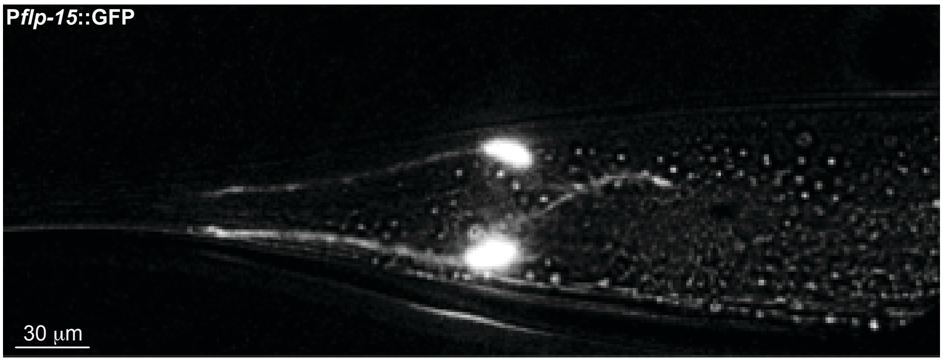

**Figure S4**

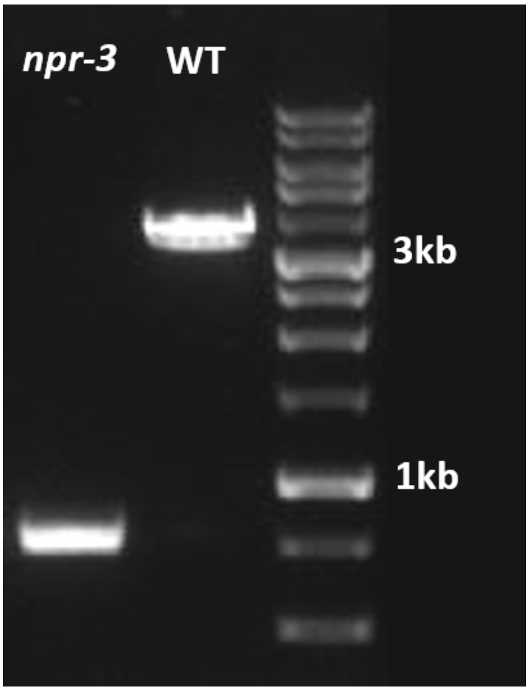

**Figure S6**

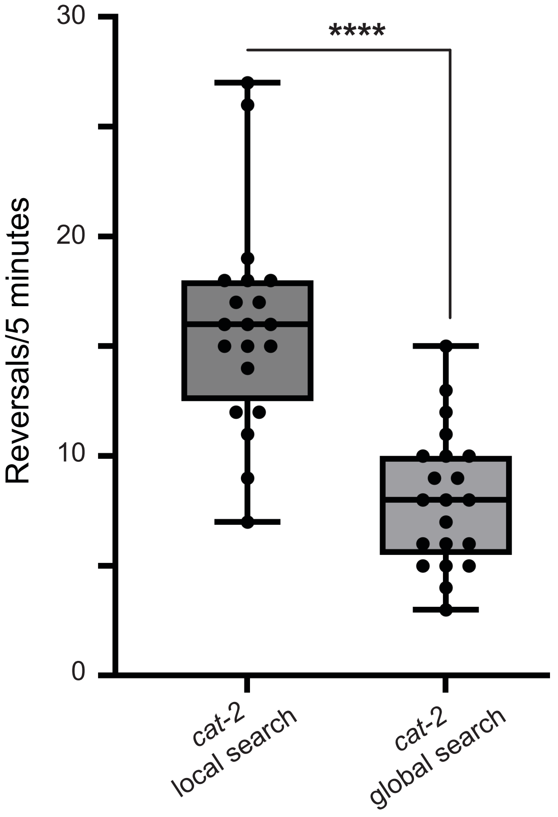

30
20
10
0
Reversals/5 minutes
****
cat-2
local search
cat-2
global search
